# Generation of an in vitro 3D multicellular culture model of ovarian high-grade serous carcinoma

**DOI:** 10.1101/2025.11.07.686515

**Authors:** Krister Wennerberg, Daria Bulanova, Laura Gall-Mas, Wojciech Senkowski, Lidia Moyano-Galceran

**Author notes:** Technical contact (Lidia Moyano-Galceran).

## Abstract

The development of translational ovarian cancer models to investigate and overcome treatment resistance accounting for the impact of the tumor microenvironment is critical. Here, we present a protocol to establish a multicellular culture model that retains both genetic complexity and the microenvironment of patient tumors, is amenable for molecular and phenotypic analyses, and high throughput drug testing. We describe steps for culturing and characterizing stromal cells derived from cryopreserved and fresh samples and detail procedures for combining them with organoids.

**Graphical abstract:** 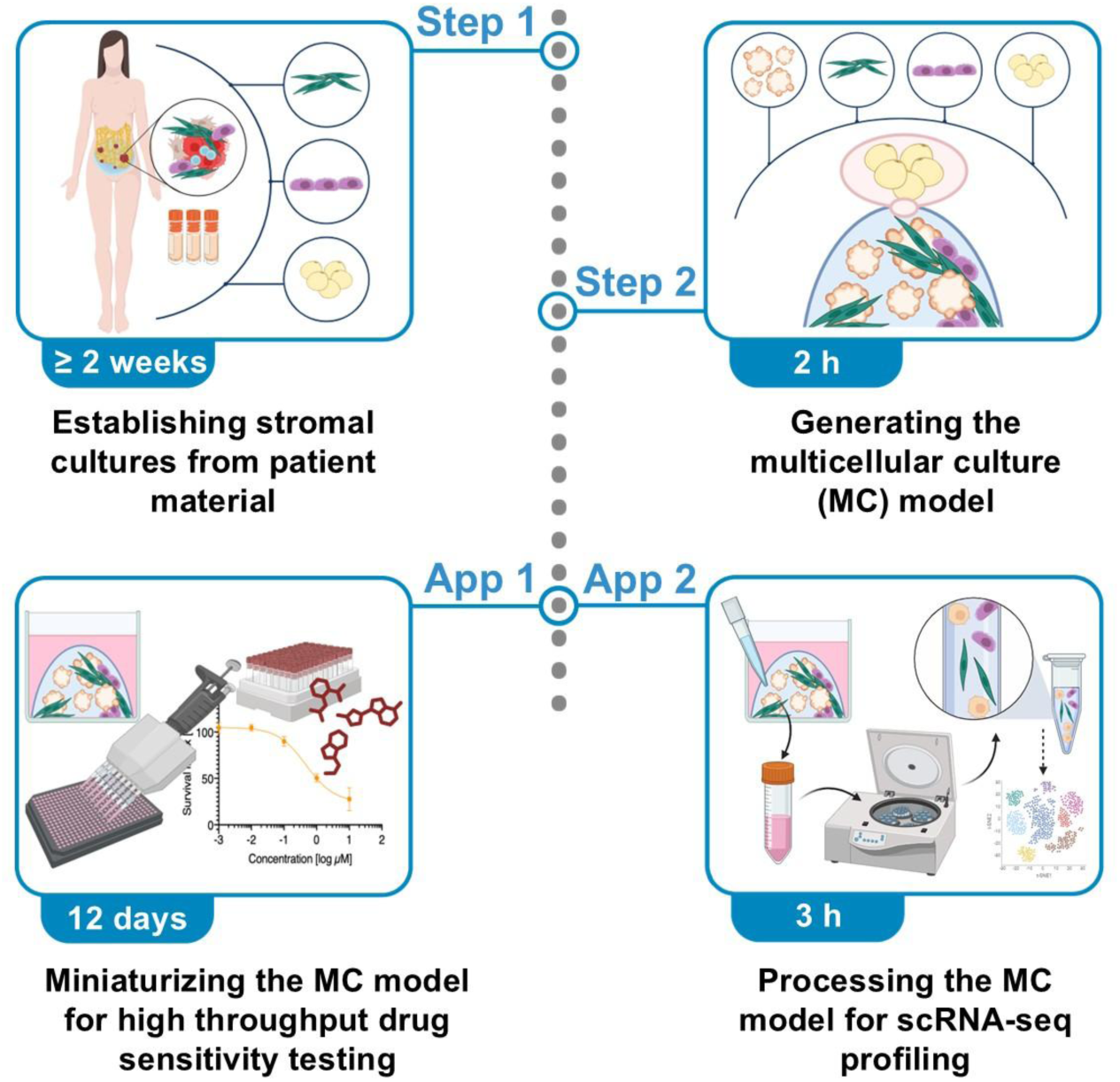

**Before you begin:** This protocol describes the *in vitro* generation of a complex 3D multicellular model (MC), mimicking relevant cellular and extracellular matrix (ECM) interactions in metastatic ovarian high-grade serous carcinoma (HGSC). First, cultures of stromal cells (cancer-associated fibroblasts (CAF), mesothelial cells and adipocytes) are generated. CAF and mesothelial cell cultures are established from fresh tumor tissues and/or from cryopreserved tissue digest and ascites fluid. Next, the identity of the stromal cells is evaluated using relevant markers, and the validated cultures are expanded and cryopreserved. Adipocytes are isolated from fresh tumor tissues and cultured in suspension for a short period before 3D embedding. Finally, previously established patient-derived cancer organoids^1^ are combined with relevant components of the tumor microenvironment (TME)^2^, including Type I collagen (main ECM protein in omental metastases) and stromal cells (**Figure 1**). The resulting MC model, which is viable for at least 14 days, can be used in various downstream applications. Here, we provide detailed protocols for two of them: high throughput drug sensitivity testing and single-cell RNA sequencing (scRNA-seq).

Figure 1.
Overview of the samples and culture conditions used to establish stromal cell cultures, and their integration with patient-derived organoids to generate the MC model.

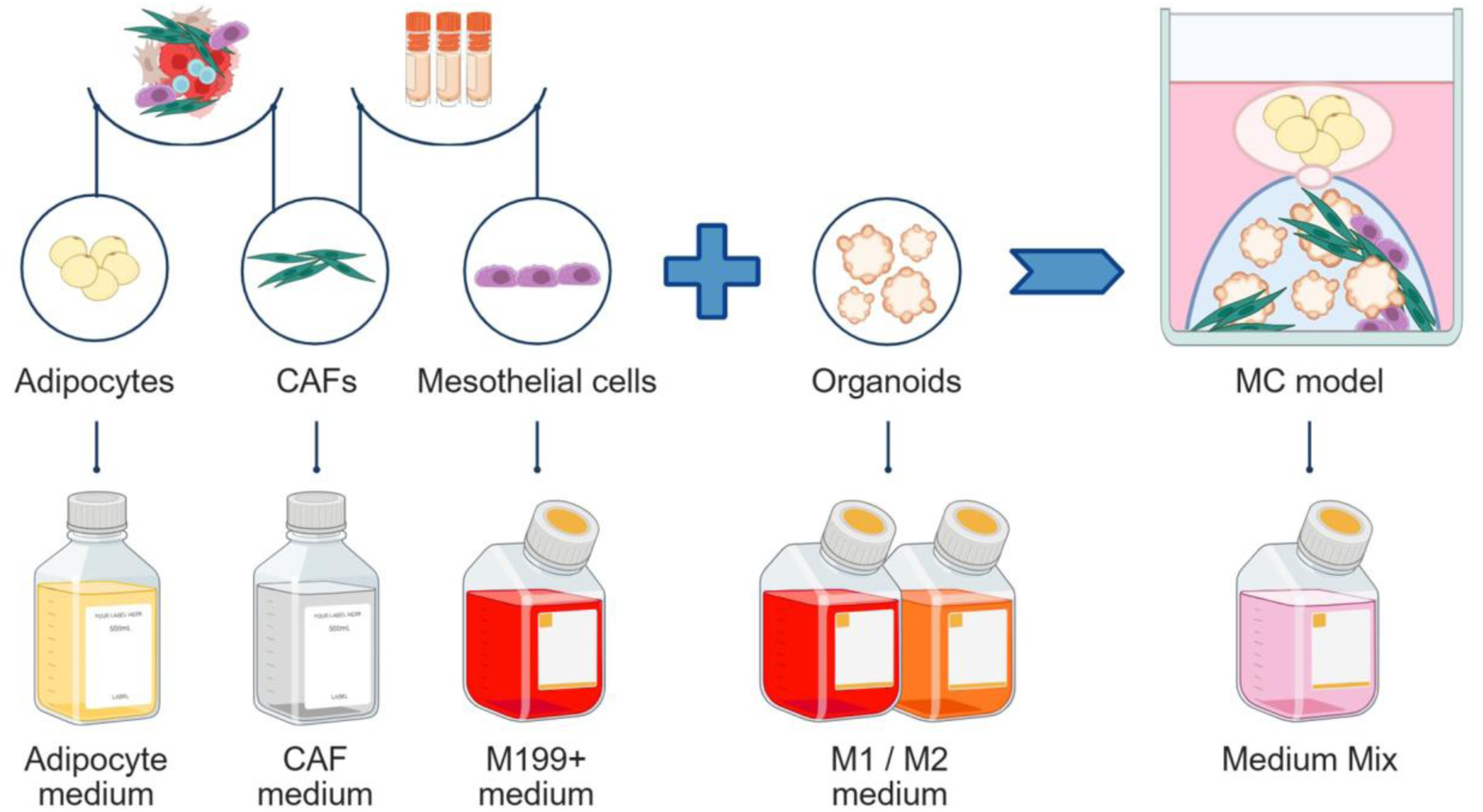

## Innovation

Compared to existing experimental *in vitro* models of HGSC, the novelty of this protocol comes from integrating patient-derived organoids with TME components, and from using cryopreserved patient material to generate stromal cell cultures. The unique MC model described here can be perturbed at the cell level (gene editing of specific cell types before assembly) and at the full model level (drug treatment), it can be studied at the single-cell level (scRNA-seq) and it can be miniaturized to perform high throughput drug sensitivity testing. In addition, the described methodology is likely adaptable for the generation of complex experimental models of other solid tumors.

### Institutional permissions

Researchers intending to replicate this protocol must ensure that they obtain the necessary permissions and approvals from their respective institutional committees. For our study, the use of all clinical materials was approved by The Ethics Committee of the Hospital District of Southwest Finland (EMTK: 145/1801/2015) and The Swedish Ethical Review Agency (2019-05149 and 2015/1862-32). All patients signed an informed consent for the use of the samples in research.

### Culture of previously established patient-derived organoids

#### Timing: 20-28 days

In this step, previously established patient-derived organoids (as described in Senkowski *et al*.^1^) are thawed, cultured and passaged until they are actively proliferating and ready to use for generating the MC model. Each organoid culture grows in a specific medium (M1 or M2; see <u>Materials and equipment</u>), which is determined during the generation of the cultures.

1. Thaw a vial of Cultrex and dilute the required amount to 7.5 mg/mL with cold PBS. **CRITICAL**: Avoid complete thawing of Cultrex at room temperature or it will polymerize; place it on ice and work on ice.
2. Thaw the cryopreserved organoids in a water bath at 37°C by swirling the cryovial.
3. Transfer the cell suspension into a tube containing 5 mL of pre-warmed M1 with 5 µM Y-27632.
4. Centrifuge at 300 x g for 5 min, aspirate the supernatant.
5. Resuspend the cell pellet in the appropriate volume of 7.5 mg/mL Cultrex (200 µL to seed 1 well of a 6-well plate).
6. Seed 10 domes of 20 µL per well in a pre-warmed 6-well plate.
7. Incubate at 37°C for 45 min.
8. Add 3 mL of pre-warmed M1/M2 containing 5 µM Y-27632 per well as required by the culture.
9. Incubate the cultures for 10-14 days while changing the medium every 2-3 days (without Y-27632). **Note:** Initially after thawing, organoids may display slower growth rates; thus, continuous monitoring is required to assess the correct passaging time.
10. Wash the domes with 2 mL of pre-warmed PBS.
11. Add 2 mL of TrypLE per well, scrape the domes with a cell lifter, and mechanically digest the domes by pipetting up and down. **CRITICAL:** Dip prime the pipette tip in TrypLE to minimize cell loss.
12. Incubate the plate for 15 min at 37°C.
13. Collect the cell suspension into a tube, wash the well with 1 mL of PBS and collect in the same tube. **CRITICAL:** Dip prime the pipette tip in TrypLE to minimize cell loss.
14. Repeat steps 4-9. **Note:** Most organoid cultures are actively proliferating and regain their standard growth rate after one passage. However, if a culture does not recover, we recommend postponing the experiment to avoid generating an inadequate MC model.

### Generation of mCherry-labelled patient-derived organoids (OPTIONAL)

#### Timing: minimum 21 days

In this step, lentiviral supernatant is produced, patient-derived organoids are infected and selected, cultured and passaged until they have recovered. The mCherry-labelled organoids can be used to generate the MC model for applications that may require the distinction between cancer and stromal cells, such as high throughput drug sensitivity testing with high content confocal imaging readout.

##### Production of lentiviral supernatant

Day 1: Seed 1×10^6^ HEK293FT cells per well in a 6-well plate. Incubate O/N in DMEM high glucose with 1% Penicillin-Streptomycin (PenStrep).

Day 2: Transfection.

`o` Aspirate medium and carefully add 1 mL per well of DMEM high glucose without PenStrep.
`o` For each well, prepare 200 μL of the DNA mixture (see <u>Materials and equipment</u>) in a 1.5 mL tube.
`o` Prepare the Lipofectamine mixture in a 1.5 mL tube. For each well, mix 16 μL of Lipofectamine 2000 with 200 μL of Opti-MEM. Incubate for 3 min at room temperature (RT) and add to the DNA mixture dropwise. Invert the tube a couple of times and incubate for 20 min at RT.
`o` Add 400 μL of transfection mixture per well dropwise and incubate O/N.

Day 3: Aspirate the medium and carefully add 0.75 mL per well of DMEM high glucose with 1% PenStrep.

Day 4: Collect the medium from each well in a 50 mL tube and add 0.75 mL per well of DMEM high glucose with PenStrep. Keep the tube with the viral supernatant at 4°C.

Day 5: Collect the medium from each well in the same 50 mL tube. Centrifuge the tube at 500 x g for 5 min and filter the supernatant using a 0.45 μm cell strainer to remove potential HEK293FT cells. Aliquot and freeze at −80°C for long-term storage.

**Note**: Titrate the lentiviral supernatant following standard protocols such as Gill *et al*^3^. *Lentiviral infection and selection of infected organoids*

Day 1: Infection in 2D.

`o` Coat two wells of a 6-well plate by adding a thin layer of 70% Cultrex diluted in PBS. Incubate at 37°C for at least 1 h.
`o` Digest 1 well of patient-derived organoids as described in steps 10-13 (<u>Culture of organoids</u>), increasing the incubation time in step 12 to 25 min to further dissociate the organoids.
`o` Centrifuge the tube at 300 x g for 5 min, aspirate the supernatant and resuspend the cell pellet in 2 mL of pre-warmed M1/M2 containing 5 µM Y-27632. Filter the cell suspension using a 0.7 μm cell strainer and take a 10 µL aliquot for cell counting.
`o` Seed 3×10^5^ cells per well in a final volume of 1.5 mL containing the lentiviral supernatant (volume adjusted based on the viral titer), M1/M2 supplemented with 5 µM Y-27632 and 8 µg/mL polybrene.

**Note**: Seed an extra well without viral supernatant and use it as control during the selection process.

Day 2: Seeding infected organoids in 3D.

`o` For each well: collect the cells in a tube, wash with 1 mL of cold PBS, add 1 mL of pre-warmed TrypLE and incubate at 37°C for 15 min. Collect the cell suspension in the same tube, together with a wash of the well with 1 mL of cold PBS.
`o` Centrifuge the tube at 400 x g for 5 min, aspirate the supernatant and proceed with steps 5-8 (<u>Culture of organoids</u>).

Day 4: Remove the medium and add 3 mL of pre-warmed M1/M2 with 0.5 μg/mL puromycin per well. **Note**: Incubate the cultures for 8-12 days while changing the medium every 2-3 days, adding puromycin until all cells in the control well are dead (cells that were not infected with lentiviral supernatant). The selection process can last 5-12 days, depending on the culture. Afterwards, perform regular medium change until the organoids become confluent and are ready to be passaged.

### Preparation of 4.5 mg/mL Type I collagen

#### Timing: 1 week

In this step, Type I collagen powder is rehydrated to a 4.5 mg/mL gel. This step should be performed at least 1 week before generating the MC model. The resulting collagen gel can be stored at 4°C and should be used within 3-4 months.

15. Prepare 11 mL of 0.3% acetic acid solution by mixing pure acetic acid with PBS.
16. Place 50 mg of collagen powder (i.e. the whole product) in a 50 mL tube.
17. Add 11 mL of 0.3% acetic acid solution to the tube and vortex vigorously.
18. Place the tube in a device with rotating movement in a cold room for 1 week.

**CRITICAL**: Vortex the tube twice daily to ensure complete collagen rehydration into a gel.

### Coating cell culture surfaces with 100 µg/mL Type I collagen

#### Timing: 45 min

In this step, the collagen gel is diluted to 100 µg/mL and used for coating cell culture surfaces needed to establish CAF cultures. This step should be performed on the days that CAF cultures are established and expanded.

19. Prepare 10 mL of 100 µg/mL collagen by adding 110 µL of 4.5 mg/mL collagen to 10 mL of cold PBS in a tube and mix by inversion. **CRITICAL**: Work on ice. **Note**: Pipette the 4.5 mg/mL collagen slowly, as it is viscous.
20. Add the 100 µg/mL collagen solution to the cell culture surface. **Note**: Add 1 mL to a well of a 6-well plate, 2 mL to a T25 flask or 5 mL to a T75 flask.
21. Incubate at 37°C for 30 min.
22. Collect the collagen solution back into the tube. **Note**: This solution can be stored at 4°C and re-used multiple times if mixed in equal parts with freshly prepared one.
23. Wash the cell culture surface with PBS. **Note**: Once the PBS is removed, medium should be added to the coated cell culture surface immediately.

### Preparation of 2.25 mg/mL Type I collagen

#### Timing: 5-10 min

In this step, the collagen gel is diluted to 2.25 mg/mL and the pH adjusted to 7.5-8. This step should be performed right before using the diluted collagen gel.

24. Prepare the required amount of 2.25 mg/mL collagen by mixing equal parts of 4.5 mg/mL collagen to 2X MEM (see <u>Materials and equipment</u>). **CRITICAL**: Work on ice. **Note**: Pipette the 4.5 mg/mL collagen slowly, as it is viscous.
25. Adjust the pH of the 2.25 mg/mL collagen to 7.5-8 with 1 M NaOH. **CRITICAL**: Work on ice. Add small volumes of NaOH (e.g. 1 µL at a time to a total of 5 µL in 300 µL of 2.25 mg/mL collagen) and use pH paper strips to check the pH after each addition (see <u>Troubleshooting 1</u>). Use the diluted collagen immediately.

### Key resources table

| REAGENT or RESOURCE | SOURCE | IDENTIFIER |
| --- | --- | --- |
| **Antibodies** |  |  |
| Anti-PDGFRβ mouse monoclonal antibody clone D-6 (IF 1:100 dilution, WB 1:500 dilution) | Santa Cruz Biotechnology | Cat# sc-374573 |
| Anti-FAP rabbit polyclonal antibody (WB 1:1000 dilution) | Thermo Fisher Scientific | Cat# PA5-99458 |
| Anti-PAX8 rabbit polyclonal antibody (IF 1:300 dilution, WB 1:2500 dilution) | Proteintech | Cat# 10336-1-AP |
| Anti-GAPDH rabbit polyclonal antibody - Loading Control (WB 1:2500 dilution) | Abcam | Cat# ab9485 |
| Anti-Ki-67 mouse monoclonal antibody clone 8D5 (IF 1:800 dilution) | Cell Signaling Technology | Cat# 9449 |
| Anti-CALB2 rabbit polyclonal antibody (IF 1:1000 dilution) | Merck | Cat# HPA007305-100UL |
| Phalloidin Labeling Probes, Alexa Fluor Plus 647 (IF 1:1000 dilution) | Thermo Fisher Scientific | Cat# A30107 |
| Donkey anti-Rabbit IgG (H+L) Highly Cross-Adsorbed Secondary Antibody, Alexa Fluor 488 (IF 1:750 dilution) | Thermo Fisher Scientific | Cat# A-21206 |
| Donkey anti-Mouse IgG (H+L) Highly Cross-Adsorbed Secondary Antibody, Alexa Fluor 647 (IF 1:750 dilution) | Thermo Fisher Scientific | Cat# A-31571 |
| Goat Anti-Rabbit IgG Antibody (H+L), Peroxidase (WB 1:10 000 dilution) | Vector Laboratories | Cat# PI-1000-1 |
| Horse Anti-Mouse IgG Antibody (H+L), Peroxidase (WB 1:10 000 dilution) | Vector Laboratories | Cat# PI-2000-1 |
| **Biological samples** |  |  |
| HGSC cryopreserved samples | Turku University Hospital | N/A |
| HGSC fresh samples | Karolinska University Hospital | N/A |
| **Chemicals, peptides, and recombinant proteins** |  |  |
| Cultrex Reduced Growth Factor Basement Membrane Extract, Type 2, Pathclear | R&D systems | Cat# 3533-005-02 |
| Advanced DMEM/F-12 (1X) | Thermo Fisher Scientific | Cat# 12634010 |
| Primocin | InvivoGen | Cat# ant-pm-1 |
| HEPES (1 M) | Thermo Fisher Scientific | Cat# 15630080 |
| GlutaMAX (100X) | Thermo Fisher Scientific | Cat# 35050061 |
| N-acetyl-L-cysteine | Merck | Cat# A7250 |
| Nicotinamide | Merck | Cat# N0636 |
| B-27 Supplement (50X), serum free | Thermo Fisher Scientific | Cat# 17504044 |
| 17β-Estradiol | Merck | Cat# E8875 |
| SB202190 | MedChemExpress | Cat# HY-10295 |
| A83-01 | Merck | Cat# SML0788 |
| Recombinant human FGF-4 | Thermo Fisher Scientific | Cat# 100-31 |
| Recombinant human FGF-10 | Thermo Fisher Scientific | Cat# 100-26 |
| Recombinant human heregulin 1-β (Neuregulin-1) | Thermo Fisher Scientific | Cat# 100-03 |
| Animal-free recombinant human EGF | Thermo Fisher Scientific | Cat# AF-100-15 |
| Forskolin | MedChemExpress | Cat# HY-15371 |
| Hydrocortisone | Merck | Cat# H0888 |
| Y-27632 | MedChemExpress | Cat# HY-10071 |
| TrypLE Express Enzyme (1X), no phenol red | Thermo Fisher Scientific | Cat# 12604013 |
| Rat Collagen Type I | Merck | Cat# C7661 |
| 10X MEM | Thermo Fisher Scientific | Cat# 11430030 |
| Medium 199, Earle's Salts | Thermo Fisher Scientific | Cat# 31150022 |
| Fetal Bovine Serum | Thermo Fisher Scientific | Cat# A5256801 |
| Penicillin-Streptomycin | Thermo Fisher Scientific | Cat# 15140122 |
| Insulin-Transferrin-Selenium (ITS -G) (100X) | Merck | Cat# I3146-5ML |
| Trace elements B (1000X) | Corning | Cat# 25-022-CI |
| 10X Collagenase/hyaluronidase in DMEM | STEMCELL Technologies | Cat# 07912 |
| Fibroblast Basal Medium | ATCC | Cat# PCS-201-030 |
| Fibroblast Growth Kit-Low serum | ATCC | Cat# PCS-201-041 |
| Trypsin-EDTA (0.25%), phenol red | Thermo Fisher Scientific | Cat# 25200056 |
| CryoStor cell cryopreservation media | Merck | Cat# C2874 |
| StemPro Adipogenesis Differentiation Kit | Thermo Fisher Scientific | Cat# A1007001 |
| Collagenase from Clostridium histolyticum | Merck | Cat# C0130-100mg |
| DPBS, no calcium, no magnesium (1000 mL) | Thermo Fisher Scientific | Cat# 14190136 |
| Hoechst 33342 | Thermo Fisher Scientific | Cat# H1399 |
| DMEM, high glucose, GlutaMAX Supplement, pyruvate (for generating mCherry labelled organoids - OPTIONAL) | Thermo Fisher Scientific | Cat# 31966021 |
| Opti-MEM Reduced Serum Medium<br>(for generating mCherry labelled organoids - OPTIONAL) | Thermo Fisher Scientific | Cat# 31985062 |
| Lipofectamine 2000 Transfection Reagent<br>(for generating mCherry labelled organoids - OPTIONAL) | Thermo Fisher Scientific | Cat# 11668027 |
| Polybrene Infection / Transfection Reagent<br>(for generating mCherry labelled organoids - OPTIONAL) | Merck | Cat# TR-1003-G |
| Puromycin Dihydrochloride<br>(for generating mCherry labelled organoids - OPTIONAL) | Thermo Fisher Scientific | Cat# A1113803 |
| **Critical commercial assays** |  |  |
| CellTox Green Cytotoxicity Assay | Promega | Cat# G8741 |
| CellTiter-Glo 2.0 Cell Viability Assay | Promega | Cat# G9421 |
| **Experimental models: Cell lines** |  |  |
| Previously established patient-derived organoids | Senkowski *et al.*<sup>1</sup> | N/A |
| HEK293FT | Thermo Fisher Scientific | Cat# R70007 |
| OVCAR3 | NCI Developmental Therapeutics Program | Cat# OVCAR-3 |
| OVCAR8 | NCI Developmental Therapeutics Program | Cat# OVCAR-8 |
| **Recombinant DNA** |  |  |
| psPAX2 packaging plasmid<br>(for generating mCherry labelled organoids - OPTIONAL) | Trono Lab | Cat#12260<br>(Addgene) |
| pMD2.G envelope plasmid<br>(for generating mCherry labelled organoids - OPTIONAL) | Trono Lab | Cat# #12259<br>(Addgene) |
| pLEX307_SmaI-T2A-mCherry transfer plasmid<br>(for generating mCherry labelled organoids - OPTIONAL) | BiOrigin ApS | N/A |
| **Software and algorithms** |  |  |
| MetaXpress (image analysis) | Molecular Devices | N/A |
| SoftMax Pro 6 (data analysis) | Molecular Devices | N/A |
| GraphPad Prism 10 (data representation) | Dotmatics | N/A |
| **Other** |  |  |
| Nunc Cell-Culture Treated 6-well plates | Thermo Fisher Scientific | Cat# 140685 |
| Corning cell lifter | Merck | Cat# CLS3008 |
| Nunc Lab-Tek II Chamber Slide System; 8-well Chamber Slide w/ removable wells | Thermo Fisher Scientific | Cat# 154534PK |
| Corning CoolCell LX Cell Freezing Container | Merck | Cat# CLS432001 |
| Corning 384-well Black/Clear Bottom Low Flange Ultra-Low Attachment Microplate | Corning | Cat# 4588 |
| Reagent Reservoir 10 mL | Merck | Cat# HS120639-100EA |

### Materials and equipment

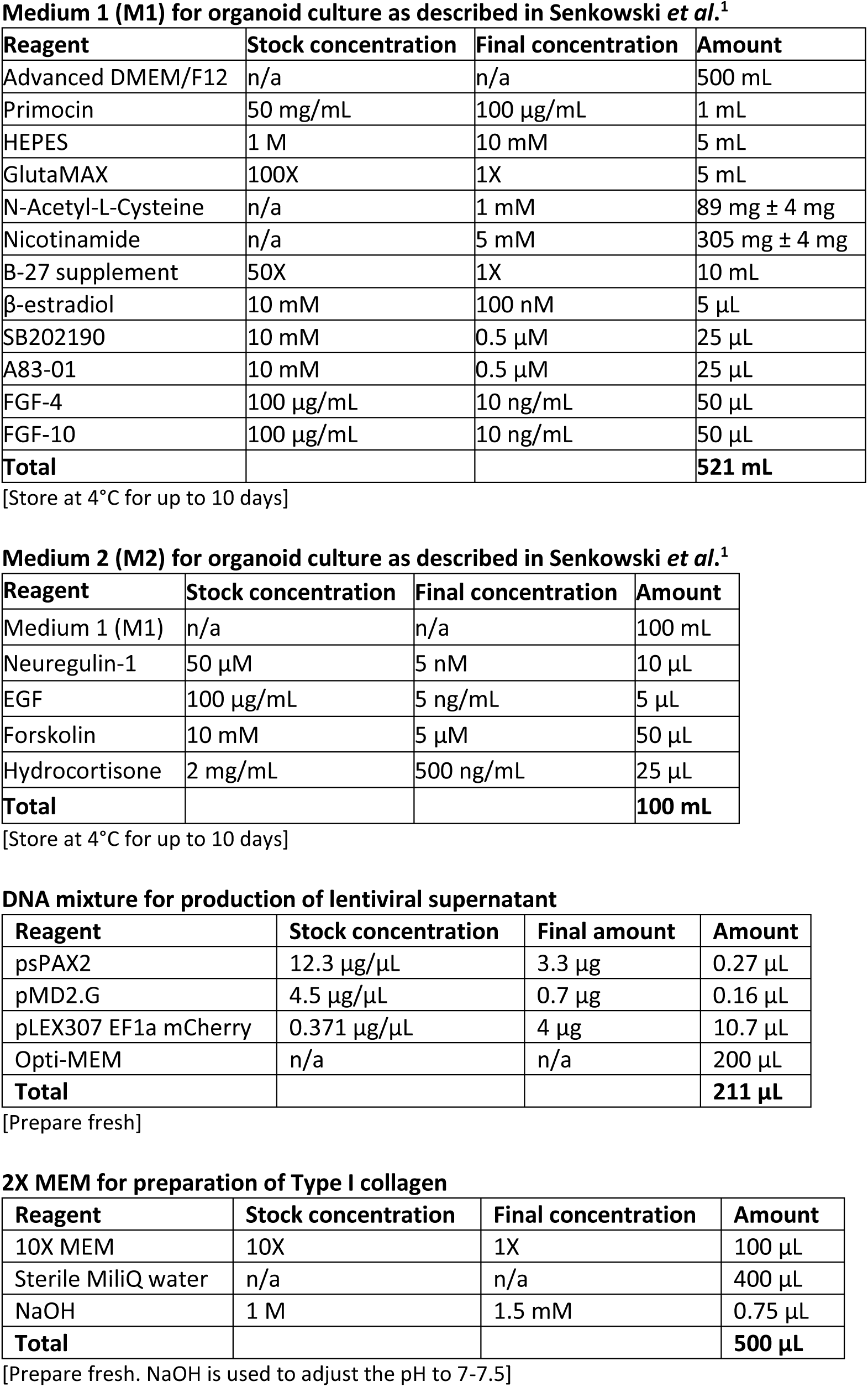
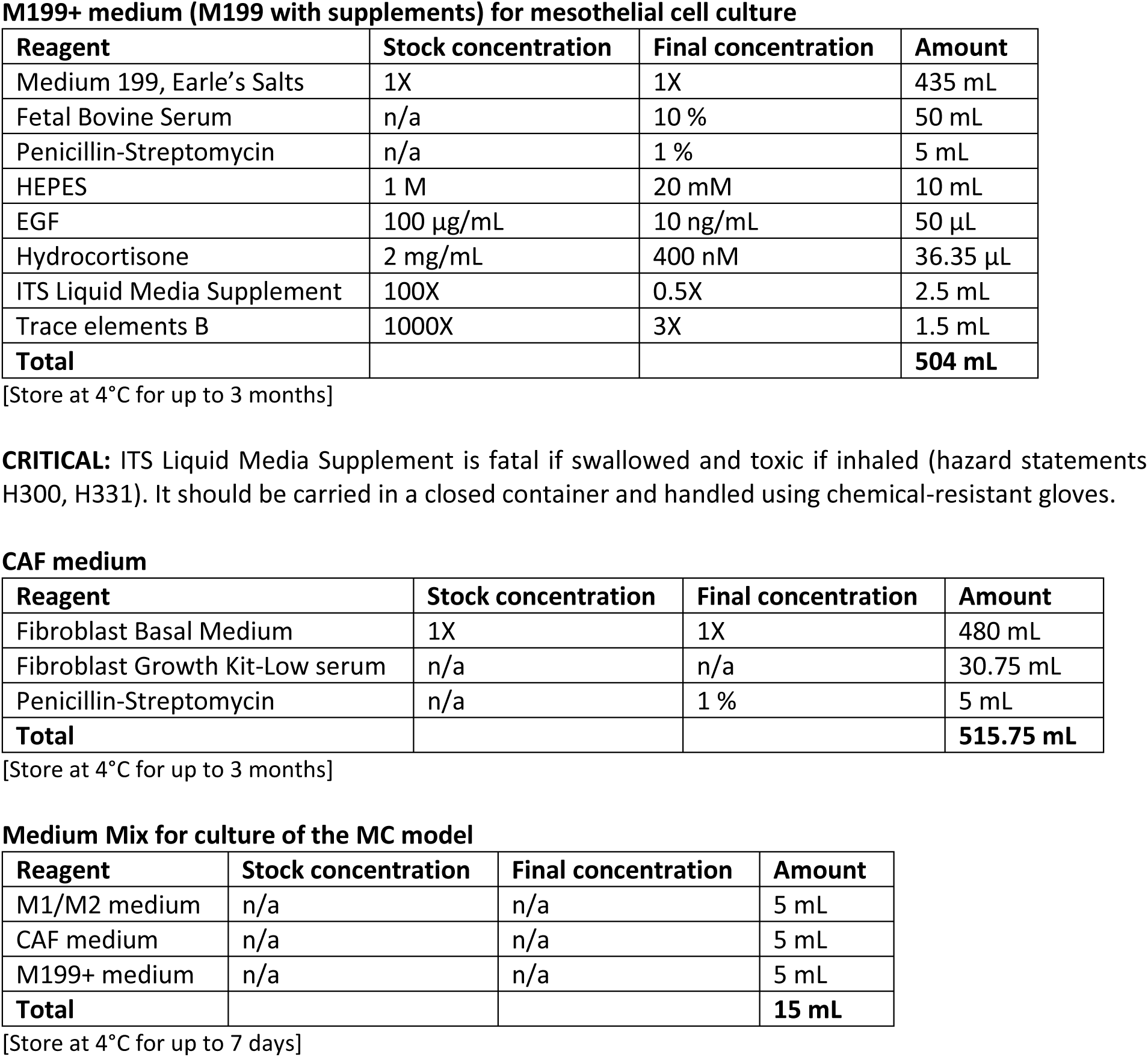

Materials and equipment Medium 1 (M1) for organoid culture as described in Senkowski *et al.*<sup>1</sup>
| Reagent | Stock concentration | Final concentration | Amount |
| --- | --- | --- | --- |
| Advanced DMEM/F12 | n/a | n/a | 500 mL |
| Primocin | 50 mg/mL | 100 µg/mL | 1 mL |
| HEPES | 1 M | 10 mM | 5 mL |
| GlutaMAX | 100X | 1X | 5 mL |
| N-Acetyl-L-Cysteine | n/a | 1 mM | 89 mg ± 4 mg |
| Nicotinamide | n/a | 5 mM | 305 mg ± 4 mg |
| B-27 supplement | 50X | 1X | 10 mL |
| β-estradiol | 10 mM | 100 nM | 5 µL |
| SB202190 | 10 mM | 0.5 µM | 25 µL |
| A83-01 | 10 mM | 0.5 µM | 25 µL |
| FGF-4 | 100 µg/mL | 10 ng/mL | 50 µL |
| FGF-10 | 100 µg/mL | 10 ng/mL | 50 µL |
| **Total** |  |  | **521 mL** |

Medium 2 (M2) for organoid culture as described in Senkowski *et al.*<sup>1</sup>
| Reagent | Stock concentration | Final concentration | Amount |
| --- | --- | --- | --- |
| Medium 1 (M1) | n/a | n/a | 100 mL |
| Neuregulin-1 | 50 µM | 5 nM | 10 µL |
| EGF | 100 µg/mL | 5 ng/mL | 5 µL |
| Forskolin | 10 mM | 5 µM | 50 µL |
| Hydrocortisone | 2 mg/mL | 500 ng/mL | 25 µL |
| **Total** |  |  | **100 mL** |

DNA mixture for production of lentiviral supernatant
| Reagent | Stock concentration | Final amount | Amount |
| --- | --- | --- | --- |
| psPAX2 | 12.3 µg/µL | 3.3 µg | 0.27 µL |
| pMD2.G | 4.5 µg/µL | 0.7 µg | 0.16 µL |
| pLEX307 EF1a mCherry | 0.371 µg/µL | 4 µg | 10.7 µL |
| Opti-MEM | n/a | n/a | 200 µL |
| **Total** |  |  | **211 µL** |

2X MEM for preparation of Type I collagen
| Reagent | Stock concentration | Final concentration | Amount |
| --- | --- | --- | --- |
| 10X MEM | 10X | 1X | 100 µL |
| Sterile MiliQ water | n/a | n/a | 400 µL |
| NaOH | 1 M | 1.5 mM | 0.75 µL |
| **Total** |  |  | **500 µL** |

**M199+ medium (M199 with supplements) for mesothelial cell culture**
| Reagent | Stock concentration | Final concentration | Amount |
| --- | --- | --- | --- |
| Medium 199, Earle's Salts | 1X | 1X | 435 mL |
| Fetal Bovine Serum | n/a | 10 % | 50 mL |
| Penicillin-Streptomycin | n/a | 1 % | 5 mL |
| HEPES | 1 M | 20 mM | 10 mL |
| EGF | 100 µg/mL | 10 ng/mL | 50 µL |
| Hydrocortisone | 2 mg/mL | 400 nM | 36.35 µL |
| ITS Liquid Media Supplement | 100X | 0.5X | 2.5 mL |
| Trace elements B | 1000X | 3X | 1.5 mL |
| **Total** |  |  | **504 mL** |

**CAF medium**
| Reagent | Stock concentration | Final concentration | Amount |
| --- | --- | --- | --- |
| Fibroblast Basal Medium | 1X | 1X | 480 mL |
| Fibroblast Growth Kit-Low serum | n/a | n/a | 30.75 mL |
| Penicillin-Streptomycin | n/a | 1 % | 5 mL |
| **Total** |  |  | **515.75 mL** |

**Medium Mix for culture of the MC model**
| Reagent | Stock concentration | Final concentration | Amount |
| --- | --- | --- | --- |
| M1/M2 medium | n/a | n/a | 5 mL |
| CAF medium | n/a | n/a | 5 mL |
| M199+ medium | n/a | n/a | 5 mL |
| **Total** |  |  | **15 mL** |

[Store at 4°C for up to 7 days]

## Step-by-step method details

### Handling fresh solid tumor samples

#### Timing: 3-4 h

Omental and peritoneal tissues with macroscopically visible tumors are collected at the operation room and stored in cold PBS during transport to the laboratory. In this step, the tissues are processed to obtain small fragments for establishing CAF cultures and fatty layer for isolation of adipocytes. As patient material is an invaluable resource, we should mention that these tissues can also be used to obtain other cell types not described in this protocol.

1. Using sterile tweezers, transfer the tissue to a 10 cm dish containing 5 mL of PBS. **Note**: If the tissue is big, cut a piece of approximately 5 cm x 5 cm and transfer it to a new dish with 5 mL of PBS.
2. Dissect part of the tissue into smaller fragments of approximately 2 mm x 2 mm by using scalpel and tweezers. **CRITICAL**: Focus on areas without necrosis, blood vessels or fat. Avoid tearing the tissue to prevent the release of necrotic cells, debris, blood and oil droplets. **Note**: These fragments should be immediately used to establish CAF cultures as described in <u>Establishing CAF cultures from tissue pieces</u>.
3. Dissect another part of the tissue into small fragments of approximately 5 mm x 5 mm by using scalpel and tweezers. **CRITICAL**: This time focus on areas that contain fat, excluding necrotic regions and blood vessels, and avoid tearing the tissue.
4. Transfer the tissue pieces into a 50 mL tube, centrifuge at 200 x g for 3 min and discard the supernatant.
5. Place the tissue pieces into a sterile beaker and cover with pre-warmed serum-free medium supplemented with 0.1-0.5X collagenase/hyaluronidase. **Note**: Adjust the concentration of collagenase/hyaluronidase depending on the tissue stiffness (e.g. use 0.1X for loose tissue). **Note**: Any serum-free cell culture medium can be used, such as M199 removed from the bottle to make the M199+ medium (see <u>Materials and equipment</u>).
6. Place a sterile magnetic bar in the beaker, cover with tin foil and leave it to stir at 37°C for 2-5 h or until the tissue has dissociated enough to recover cells. **Note**: If the original tumor tissue was rather small, the digestion can be performed in 2 mL Eppendorf tubes in a heat block with automatic shaking.
7. Collect the fatty layer with a pipette into a tube, centrifuge at 200 x g for 3 min and discard the top oily fraction as well as the residual medium at the bottom (see <u>Troubleshooting 2</u>). **Note**: The adipocyte fraction can now be used to establish adipocyte cultures as described in <u>Establishing adipocyte cultures</u>.

### Establishing CAF cultures from tissue pieces

#### Timing: minimum 2 weeks

This step describes the establishment, maintenance, characterization, expansion and cryopreservation of CAF cultures. All cell culture surfaces used for CAF culture are pre-coated with collagen (see <u>Coating</u> <u>cell culture surfaces</u>).

8. Add 3 mL of pre-warmed CAF medium (see <u>Materials and equipment</u>) in a well of a 6-well plate.
9. Place 10 tissue pieces from step 2 (<u>Handling fresh solid tumor samples</u>) in the well.
10. Incubate the cultures until cells reach 90% confluency (between 5-16 days) while changing the medium every 2-3 days without disturbing the tissue pieces.
11. Transfer the tissue pieces to a new well of a 6-well plate containing 3 mL of pre-warmed CAF medium. **Note**: Sometimes more cells can be obtained from the same tissue pieces; otherwise, discard the tissue pieces.
12. Wash the wells with 1 mL of pre-warmed PBS.
13. Add 0.5 mL of pre-warmed trypsin per well and incubate the plate for 5-10 min at 37°C until the cells detach.
14. Add 1 mL of pre-warmed CAF medium per well, resuspend the cells, transfer into a tube and take a 10 µL aliquot for cell counting.
15. Seed 1.5×10^5^ cells in a T25 flask or 3×10^5^ cells in a T75 flask.
16. Incubate the cultures until cells reach 90% confluency while changing the medium every 2-3 days.
17. Wash the cells with 1-2 mL of pre-warmed PBS.
18. Add 0.5 mL per T25 flask or 1.5 mL per T75 flask of pre-warmed trypsin and incubate for 5-10 min at 37°C until the cells detach.
19. Add 1 mL per T25 flask or 1.5 mL per T75 flask of pre-warmed CAF medium, resuspend the cells, transfer into a tube and take a 10 µL aliquot for cell counting.
20. Seed 1×10^5^ cells in a well of a 6-well plate. **Note**: Once confluent, characterize the culture by western blot (e.g. using the protocol by Moyano-Galceran *et al*.^4^). See <u>Key Resources Table</u>, <u>Expected outcomes</u> and <u>Troubleshooting</u> <u>4</u> for more details.
21. Repeat steps 15-19 with the remaining cells. **Note**: We recommend expanding CAF cultures and cryopreserving cells at every passage once their identity has been confirmed. **Note**: Since untransformed CAFs have limited growth potential, we recommend stopping the expansion of CAF cultures once their growth rate slows down and their morphology changes.
22. Centrifuge the tubes containing cells at 300 x g for 5 min and aspirate the supernatant.
23. Resuspend the cell pellet in 1 mL of CryoStor cell cryopreservation medium and transfer into a cryovial. **Note**: We recommend cryopreserving up to 4.5×10^5^ cells per cryovial.
24. Place the cryovial in a CoolCell freezing container and store it in the-80°C freezer shortly. **Note**: Transfer the cryovial to a liquid nitrogen tank for long-term storage.

### Establishing adipocyte cultures

#### Timing: 1 week

This step describes the establishment and maintenance of adipocyte cultures in Adipocyte medium (i.e. StemPro Adipogenesis Differentiation Kit; see <u>Key Resources Table</u>).

25. Add 4 mL of pre-warmed Adipocyte medium into a T25 flask.
26. Resuspend the adipocyte fraction from step 7 (<u>Handling fresh solid tumor samples</u>) in 1 mL of pre-warmed Adipocyte medium and transfer to the T25 flask.
27. Add 2 mL of fresh Adipocyte medium after 3-4 days and incubate for a total of 1 week.
28. Transfer the cell suspension to a tube, centrifuge at 200 x g for 5 min and aspirate the supernatant.
29. Use a pipette to estimate the volume of adipocytes by taking up the adipocyte fraction.
30. Mix the adipocyte fraction with 2.25 mg/mL collagen (see <u>Preparation of 2.25 mg/mL Type I</u> <u>collagen</u>) in a 1:4 ratio. **CRITICAL**: Work on ice and at a steady pace to avoid complete collagen polymerization.
31. Seed 10 domes of 20 µL per well in a pre-warmed 6-well plate.
32. Incubate at 37°C for 1 h.
33. Gently add 3 mL of pre-warmed Adipocyte medium per well (see <u>Troubleshooting 3</u>). **Note**: Adipocyte gels can now be used to generate the MC model. These cultures will remain viable for up to 30 days.

### Thawing cryopreserved patient samples

#### Timing: 10 min

An alternative source of stromal cells to fresh patient material can be cryopreserved tumor tissue digest and enriched cell suspensions from ascites. In this step, cryopreserved samples (obtained as described in Senkowski *et al*.^1^) are thawed before being used for establishing stromal cell cultures.

34. Thaw the cryopreserved samples in a water bath at 37°C by swirling the cryovials.
35. Transfer the cell suspensions into tubes containing 5 mL of pre-warmed serum-free medium. **Note**: Any serum-free cell culture medium can be used, such as M199 removed from the bottle to make the M199+ medium (see <u>Materials and equipment</u>).
36. Centrifuge at 300 x g for 5 min and aspirate the supernatant.
37. Resuspend the cell pellets in pre-warmed serum-free medium and take a 10 µL aliquot for cell counting.
38. Divide the cell suspensions into different tubes with 2.5×10^5^ cells/ tube.
39. Centrifuge at 300 x g for 5 min and aspirate the supernatant. **Note**: Cells are now ready for resuspension in cell type specific medium as described in <u>Establishing CAF cultures from cryopreserved patient samples</u> and <u>Establishing mesothelial</u> <u>cell cultures from cryopreserved patient samples</u>.

### Establishing CAF cultures from cryopreserved patient samples

#### Timing: minimum 2 weeks

This step describes the establishment of CAF cultures, which are then maintained, characterized, expanded and cryopreserved as described in <u>Establishing CAF cultures from tissue pieces</u>. Although CAF cultures can be established from cryopreserved enriched cell suspensions from ascites, it is often more effective to use cryopreserved tumor tissue digest.

40. Add 1 mL of pre-warmed CAF medium (see <u>Materials and equipment</u>) in a well of a 6-well plate pre-coated with collagen (see <u>Coating cell culture surfaces</u>).
41. Resuspend the cell pellet from step 39 (<u>Thawing cryopreserved patient samples</u>) in 1 mL of pre-warmed CAF medium and transfer to the well.
42. Incubate the culture until cells reach 90% confluency while changing the medium every 2-3 days.
43. Follow steps 12-24 (Establishing CAF cultures from tissue pieces).

### Establishing mesothelial cell cultures from cryopreserved patient samples

#### Timing: minimum 2 weeks

This step describes the establishment, maintenance, characterization, expansion and cryopreservation of mesothelial cell cultures. Although mesothelial cell cultures can be established from cryopreserved tumor tissue digest, it is often more effective to use cryopreserved enriched cell suspensions from ascites.

44. Resuspend the cell pellet from step 39 (<u>Thawing cryopreserved patient samples</u>) in 5 mL of pre-warmed M199+ medium (see <u>Materials and equipment</u>) and transfer to a T25 flask.
45. Incubate the flask until cells reach 90% confluency while changing the medium every 2-3 days.
46. Wash the cells with 1 mL of pre-warmed PBS.
47. Add 0.5 mL of pre-warmed trypsin and incubate the flask for 5-10 min at 37°C until the cells detach.
48. Add 1 mL of pre-warmed M199+ medium per T25 flask, resuspend the cells, transfer into a tube and take a 10 µL aliquot for cell counting.
49. Seed 1.5×10^5^ cells in a T25 flask or 3×10^5^ cells in a T75 flask.
50. Incubate the cultures until cells reach 90% confluency while changing the medium every 2-3 days.
51. Wash the cells with 1-2 mL of pre-warmed PBS.
52. Add 0.5 mL per T25 flask or 1.5 mL per T75 flask of pre-warmed trypsin and incubate for 5-10 min at 37°C until the cells detach.
53. Add 1 mL per T25 flask or 1.5 mL per T75 flask of pre-warmed M199+ medium, resuspend the cells, transfer into a tube and take a 10 µL aliquot for cell counting.
54. Seed 1.5×10^4^ cells per well in 4 wells of an 8-well chamber slide. **Note**: Once confluent, use these cells to characterize the culture by immunofluorescence staining (e.g. using the protocol by Moyano-Galceran *et al*.^4^). See <u>Key Resources Table</u>, <u>Expected outcomes</u> and <u>Troubleshooting 4</u> for more details.
55. Repeat steps 49-53 with the remaining cells. **Note**: We recommend expanding the cultures and cryopreserving cells at every passage once their identity has been confirmed. **Note**: Since untransformed mesothelial cells have limited growth potential, we recommend stopping the expansion of the cultures once their growth rate slows down and their morphology changes.
56. Centrifuge the tubes containing cells at 300 x g for 5 min and aspirate the supernatant.
57. Resuspend the cell pellet in 1 mL of CryoStor cell cryopreservation medium and transfer into a cryovial. **Note**: We recommend cryopreserving up to 4.5×10^5^ cells per cryovial.
58. Place the cryovial in a CoolCell freezing container and store it in the-80°C freezer shortly. **Note**: Transfer the cryovial to a liquid nitrogen tank for long-term storage.

***Pause: All the cell types required to generate the MC model are now ready.***

### Generating the MC model

#### Timing: 2 h

In this step, the patient-derived organoids that have recovered from the cryopreservation process (see <u>Culture of organoids</u>) and have grown to the time of passage are combined with actively growing CAFs and mesothelial cells by embedding them in a mixture of Cultrex and Type I collagen. In this protocol, the MC model contains 15% of stromal cells, of which 12.5% are CAFs and 2.5% are mesothelial cells, recreating HGSC tumors^5^. In addition, adipocyte gels can be attached to the MC model to better recapitulate the metastatic fatty omentum. The MC model is typically established in 6-well plates and grown for up to 14 days.

59. Thaw a vial of Cultrex and dilute the required amount to 7.5 mg/mL with cold PBS. **CRITICAL**: Avoid complete thawing of Cultrex at room temperature or it will polymerize; place it on ice and work on ice.
60. Trypsinize and count stromal cells:

a. Wash the cells with 1-2 mL of pre-warmed PBS.
b. Add 0.5 mL per T25 flask or 1.5 mL per T75 flask of pre-warmed trypsin and incubate for 5-10 min at 37°C until the cells detach.
c. Add 1 mL per T25 flask or 1.5 mL per T75 flask of pre-warmed cell type specific medium, resuspend the cells, transfer into tubes and take 10 µL aliquots for cell counting.
61. Digest 1 well of patient-derived organoids:

a. Wash the domes with 2 mL of pre-warmed PBS.
b. Add 2 mL of TrypLE, scrape the domes with a cell lifter, and mechanically digest the domes by pipetting up and down. **CRITICAL**: Dip prime the pipette tip in TrypLE to minimize cell loss.
c. Incubate the plate for 15 min at 37°C.
d. Collect the cell suspension into a tube, wash the well with 1 mL of PBS and collect in the same tube. **CRITICAL**: Dip prime the pipette tip in TrypLE to minimize cell loss.
62. Prepare the cell mixture:

a. Add 1.8×10^5^ CAFs and 3.6 x10^4^ mesothelial cells into the tube containing the organoid cell suspension.
b. Centrifuge at 300 x g for 5 min and aspirate the supernatant.
63. Prepare the mixture of matrices (see <u>Troubleshooting 5</u>):

a. Pipette the required amount of 4.5 mg/mL collagen (see <u>Preparation of 4.5 mg/mL</u> <u>Type I collagen</u>) into a tube. **CRITICAL**: Work on ice and pipette slowly as collagen is viscous. **Note**: Collagen and Cultrex are mixed in 1:1 ratio to generate the MC model.
b. Adjust the pH of the 4.5 mg/mL collagen to 7.5-8 with 1 M NaOH. **CRITICAL**: Work on ice and pipette slowly as collagen is viscous. Add small volumes of NaOH (e.g. 5-10 µL at a time to a total of 40 µL in 1 mL of 4.5 mg/mL collagen) and use pH paper strips to check the pH after each addition (see <u>Troubleshooting 1</u>).
c. Add the required amount of 7.5 mg/mL Cultrex and mix by pipetting gently. **CRITICAL**: Work on ice and pipette slowly as the mixture of matrices is viscous. Avoid forming bubbles.
64. Resuspend the cell pellet in the appropriate volume of the mixture of matrices (200 µL to seed 1 well of a 6-well plate).
65. Seed 10 domes of 20 µL per well in a pre-warmed 6-well plate.
66. Incubate at 37°C for 1 h. **Optional**: Adipocyte gels can be added to the MC model after 30 min incubation as follows: Place a 3 µL droplet of 4.5 mg/mL collagen on top of a dome. Carefully take up an adipocyte gel (see steps 25-33 of <u>Establishing adipocyte cultures</u>) with a pipette tip and place it on top of the small collagen droplet. Repeat this step as many times as desired. Alternatively, place the adipocyte gels in between domes and use 3 µL droplets of 4.5 mg/mL collagen to attach them. Incubate at 37°C for 30 min.
67. Add 3 mL of pre-warmed M1/M2-based Medium Mix (see <u>Materials and equipment</u>) containing 5 µM Y-27632 per well as required by the culture.
68. Incubate the cultures for 10-14 days while changing the medium every 2-3 days (without Y-27632).

### Miniaturizing the MC model for high throughput drug sensitivity testing

#### Timing: 12 days

In this step, the MC model is seeded in a 384-well plate and cultured for 12 days with several medium changes. On day 7, the MC model is exposed to a 48-h drug treatment, and on day 12, the cell viability/cytotoxicity endpoint readout is conducted. We perform medium aspiration with BioTek EL405 LS microplate washer (Agilent Technologies), dispense drugs with Echo 550 acoustic liquid handler (Labcyte), and acquire high content confocal microscope images with ImageXpress Confocal HT.ai imaging system (Molecular Devices) and/or luminescent signal using a SpectraMax Paradigm Multi-Mode Detection Platform (Molecular Devices).

69. Place a 384-well plate and a 10 mL sterile reservoir at-20°C. After 15 min, move the plate to 4°C until ready to perform the seeding.
70. Perform steps 59-63 (<u>Generating the MC model</u>).
71. Resuspend the cell pellet in the appropriate volume of the mixture of matrices (e.g. 4 mL to seed one 384-well plate, leaving the top and bottom rows empty) (see <u>Troubleshooting 5</u>).
72. Place the pre-chilled 10 mL sterile reservoir and 384-well plate on ice.
73. Transfer the resuspended cell pellet to the reservoir and seed 10 µL per well with a multichannel electronic pipette (e.g. 8 Channel VIAFLO Electronic Pipette 10 – 300 µL, Integra Biosciences). **CRITICAL**: Set the aspiration and dispensing speeds of the multichannel electronic pipette to lowest to facilitate working with the viscous mixture of matrices.
74. Incubate on ice for 15 min and subsequently at 37°C for 30 min.
75. Using a multichannel electronic pipette, add 40 µL of pre-warmed M1/M2-based Medium Mix (see <u>Materials and equipment</u>) containing 5 µM Y-27632 per well as required by the culture. Centrifuge the plate at 300 x g for 15 sec. **CRITICAL**: Set the dispensing speed of the multichannel electronic pipette to lowest to minimize matrix disruption.
76. Incubate the cultures for 3 days.
77. Perform medium change by aspirating 30 µL of medium from each well with a microplate washer (e.g. BioTek EL405 LS microplate washer, Agilent Technologies) and adding 30 µL of pre-warmed M1/M2-based Medium Mix (see <u>Materials and equipment</u>) per well as required by the culture with a multichannel electronic pipette. **CRITICAL**: Set the aspiration speed of the microplate washer and the aspiration and dispensing speeds of the multichannel electronic pipette to lowest to minimize matrix disruption.
78. Incubate the cultures for 4 days.
79. On day 7, perform medium change as described in step 77 prior to adding the drug treatment (e.g. carboplatin) with Echo 550 acoustic liquid handler (Labcyte). **Optional**: Include a positive cell death control such as 10 µM bortezomib and a vehicle control (e.g. 0.1% DMSO) if the chosen drug is not dissolved in medium or water.
80. After the desired drug exposure time (e.g. 48 h), perform medium change as described in step 77.
81. Incubate the cultures for 3 more days.
82. On day 12, aspirate 25 µL of medium from each well with a microplate washer and perform the following endpoint measurements:

a. CellTox Green Cytotoxicity Assay: **Note**: If the organoids are mCherry-labelled (see <u>Generation of mCherry-labelled</u> <u>organoids</u>), this readout allows the separation of the cell death signal coming from cancer cells (simultaneously red and green fluorescent) versus stromal cells (only green fluorescent).

i. Prepare the CellTox Green reagent by mixing 20 µL of Green Dye for each 10 mL of Assay Buffer. Add 20 µg/mL of Hoechst 33342 to the staining solution (the final concentration will be 10 µg/mL).
ii. Using a multichannel electronic pipette, add 25 µL of staining solution per well.
iii. Mix the plate by orbital shaking at 700-900 revolutions per minute (rpm) for 1 min and incubate for 2 h at RT in the dark.
iv. Capture green, red (optional) and blue fluorescence signals with ImageXpress Confocal HT.ai high-content imaging system (Molecular Devices). **Note**: Use the following excitation and emission wavelengths: 467.5/21 nm and 520/28 nm for green signal, 555 nm and 624/40 nm for red signal, and 405/20 nm and 452/45 nm for blue signal.
v. Perform image analysis.
b. CellTiter-Glo 2.0 Cell Viability Assay: **Note**: Use this assay when the organoids are not mCherry-labelled or when high content imaging readout is not needed. **Note**: CellTiter-Glo 2.0 Cell Viability Assay can also be performed after CellTox Green Cytotoxicity Assay as a multiplexed readout. In this case, aspirate again 25 µL of medium from each well with a microplate washer before proceeding.

i. Using a multichannel electronic pipette, add 25 µL of CellTiter-Glo 2.0 Reagent per well.
ii. Mix the plate by orbital shaking at 700-900 rpm for 5 min and incubate for 30 min at RT in the dark.
iii. Measure the luminescence signal emission at 578 nm with SpectraMax Paradigm Multi-Mode Detection Platform (Molecular Devices).
iv. Perform data analysis.

### Processing the MC model for scRNA-seq profiling

#### Timing: 3 h

In this step, the cells from the MC model are retrieved after the desired experiment has been performed (e.g. 12-day culture with an 8 h drug exposure on day 6) and processed prior to library preparation with the Chromium next GEM Single Cell 3’ Reagent Kit v.3.1 (10X Genomics).

83. Prepare 1 mg/mL Type I collagenase by resuspending the lyophilized collagenase enzyme in serum-free medium and vortex until dissolved. **CRITICAL:** Keep the collagenase solution on ice. **Note**: Any serum-free cell culture medium can be used, such as M199 removed from the bottle to make the M199+ medium (see <u>Materials and equipment</u>).
84. If the MC model contained adipocyte gels, remove them with a pipette tip and discard them. **Note**: Adipocytes are technically challenging cells to sequence with scRNA-seq due to their high buoyancy, fragility and large size, currently being incompatible with conventional droplet-based single-cell platforms.
85. Wash the domes with 2 mL of pre-warmed PBS per well.
86. Add 1 mL of 1 mg/mL collagenase solution per well and scrape the domes with a cell lifter.
87. Incubate the plate for 15 min at 37°C.
88. Mechanically digest the domes by pipetting up and down. Collect the cell suspension into a tube, wash the well with 1 mL of ice-cold PBS and collect in the same tube. **CRITICAL**: Dip prime the pipette tip in collagenase solution to minimize cell loss.
89. Add 8 mL of ice-cold PBS to the tube and mix by inversion.
90. Centrifuge at 300 x g for 5 min and discard the supernatant.
91. Resuspend the cell pellet in 2 mL of TrypLE. **CRITICAL**: Dip prime the pipette tip in TrypLE to minimize cell loss.
92. Incubate the tube for 30 min at 37°C.
93. Mechanically digest the domes by pipetting up and down. Add 1 mL of ice-cold PBS to the tube and mix by inversion. **CRITICAL**: Dip prime the pipette tip in TrypLE to minimize cell loss.
94. Centrifuge at 300 x g for 5 min and discard the supernatant.
95. Resuspend the cell pellet in 5 mL of ice-cold DPBS and keep on ice for 15 min. **CRITICAL**: Dip prime the pipette tip in DPBS to minimize cell loss.
96. Centrifuge at 300 x g for 5 min and discard the supernatant.
97. Resuspend the cell pellet in 5 mL of ice-cold DPBS containing 0.1 mM EDTA. **CRITICAL**: Dip prime the pipette tip in DPBS to minimize cell loss.
98. Centrifuge at 300 x g for 5 min and discard the supernatant.
99. Resuspend the cell pellet in 5 mL of ice-cold DPBS containing 0.1 mM EDTA. **CRITICAL**: Dip prime the pipette tip in DPBS to minimize cell loss.
100. Filter the cell suspension through a 40 µm cell strainer and collect into a new tube.
101. Centrifuge at 150 x g for 5 min and discard the supernatant.
102. Resuspend the cell pellet in 5 mL of ice-cold DPBS containing 0.1 mM EDTA. **CRITICAL**: Dip prime the pipette tip in DPBS to minimize cell loss.
103. Centrifuge at 150 x g for 5 min and discard the supernatant. **Note**: The cell pellet should now be resuspended in a small volume of ice-cold DPBS containing 0.1 mM EDTA to perform sample quality check (e.g. using PI-acridine orange staining and Luna-FX7 cell counter) before proceeding with library preparation.

### Expected outcomes

This protocol enables the establishment, maintenance, characterization, expansion and cryopreservation of CAF and mesothelial cell cultures from fresh and/or cryopreserved HGSC tumor tissues and tissue digest, and from cryopreserved ascites fluid. Although CAF cultures established from different patients and sources may present different growth patterns, they should exhibit a spindle-shape morphology and express PDGFRβ (may also express FAP) while simultaneously being devoid of PAX8 expression (marker of HGSC cancer cells) (**Figure 2**). Instead, mesothelial cell cultures may present similar growth rates, exhibit a cobblestone morphology and express calretinin (**Figure 3**). Both CAFs and mesothelial cells can be cryopreserved and later placed back in culture, becoming a versatile tool for different applications besides the generation of the MC model. This protocol also allows the establishment and maintenance of adipocyte cultures by embedding adipocytes in collagen gels (**Figure 4**). The adipocyte cultures cannot be cryopreserved and are viable for up to 30 days; thus, they should be used for the generation of the MC model right after establishment.

**Figure 2.**
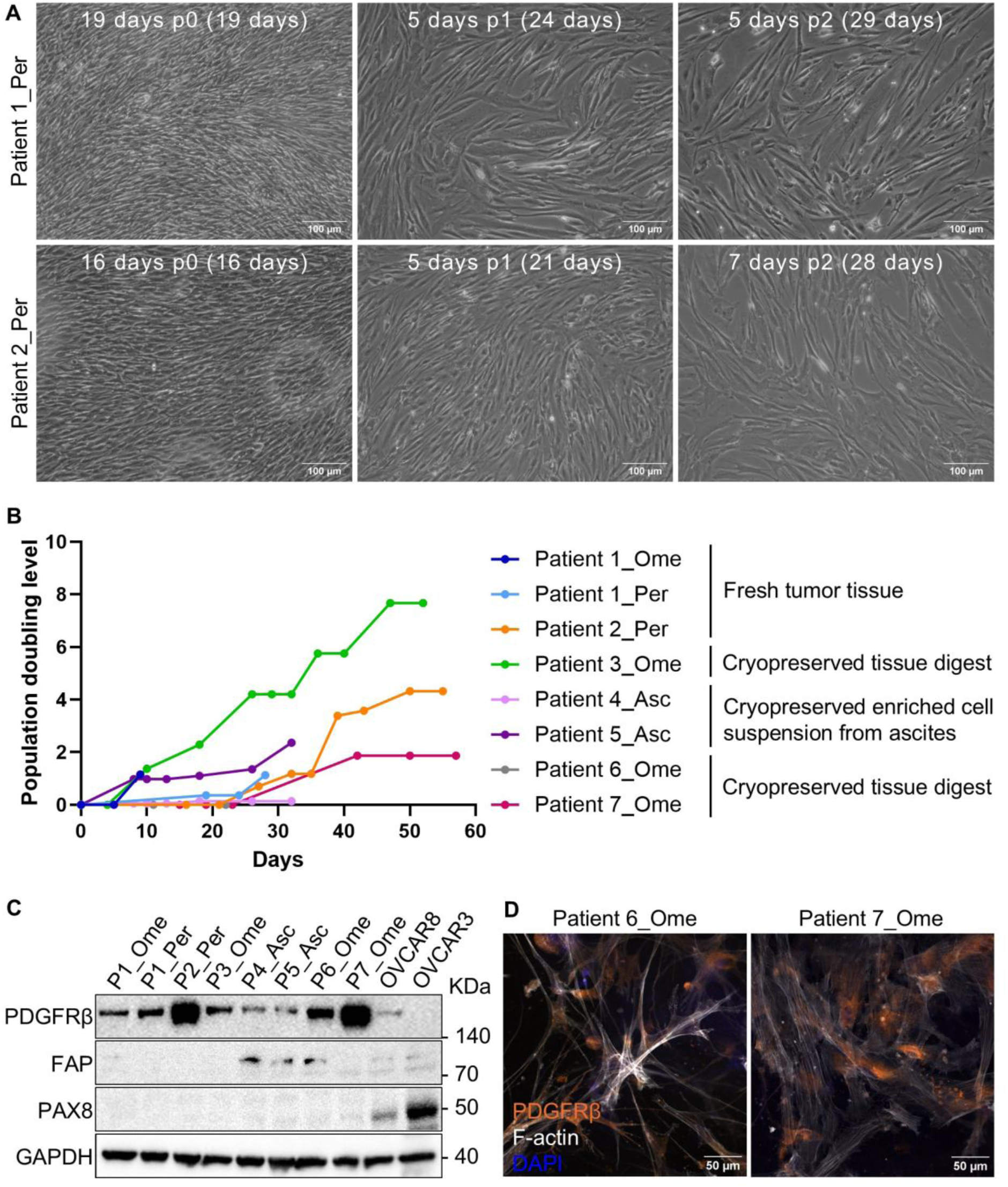
CAF cultures. **A)** Representative 10X phase-contrast images of 2 patient-derived CAF cultures displaying the characteristic spindle-shape morphology at different passages. The number of days in each passage, the passage number as well as the total number of days in culture are indicated in each image. Images were taken with a digital camera (Leica DFC340 FX) attached to an optical microscope. Scale bar: 100 µm. **B)** Chart displays the growth of CAF cultures as changes in population doubling level (PDL) over time. The type of patient material used to establish each culture is indicated and different growth dynamics can be seen. PDL at each passage was the cumulative sum of PD, which was calculated with the following formula at each passage: 𝑃𝐷 = 𝐿𝑜𝑔2((𝐹𝑖𝑛𝑎𝑙 𝑐𝑒𝑙𝑙 𝑐𝑜𝑢𝑛𝑡 − 𝐼𝑛𝑖𝑡𝑖𝑎𝑙 𝑐𝑒𝑙𝑙 𝑐𝑜𝑢𝑛𝑡)/𝐼𝑛𝑖𝑡𝑖𝑎𝑙 𝑐𝑒𝑙𝑙 𝑐𝑜𝑢𝑛𝑡) **C)** Immunoblotting for PDGFRβ and FAP (CAF markers) and PAX8 (HGSC marker) in 8 established CAF cultures and 2 HGSC cell lines (i.e. OVCAR8 and OVCAR3; used as positive controls for PAX8 expression). All CAF cultures expressed PDGFRβ while FAP was only detected in cultures derived from Patients 4-6. GAPDH was used as loading control. **D)** Representative 20X confocal fluorescence images of 2 patient-derived CAF cell cultures stained for PDGFRβ and F-actin (cytoskeleton; phalloidin probe). Images were taken with Zeiss Laser Scanning Confocal Microscope (LSM800). Scale bar: 50 µm. Abbreviations: Asc – ascites fluid, Ome – omentum, Per – peritoneum. The data that was used to generate B) is provided in Supplementary Table 1. The uncropped images from C) are provided as Supplementary Figure 1. The single channel images from D) are provided as Supplementary Figure 2.

**Figure 3.**
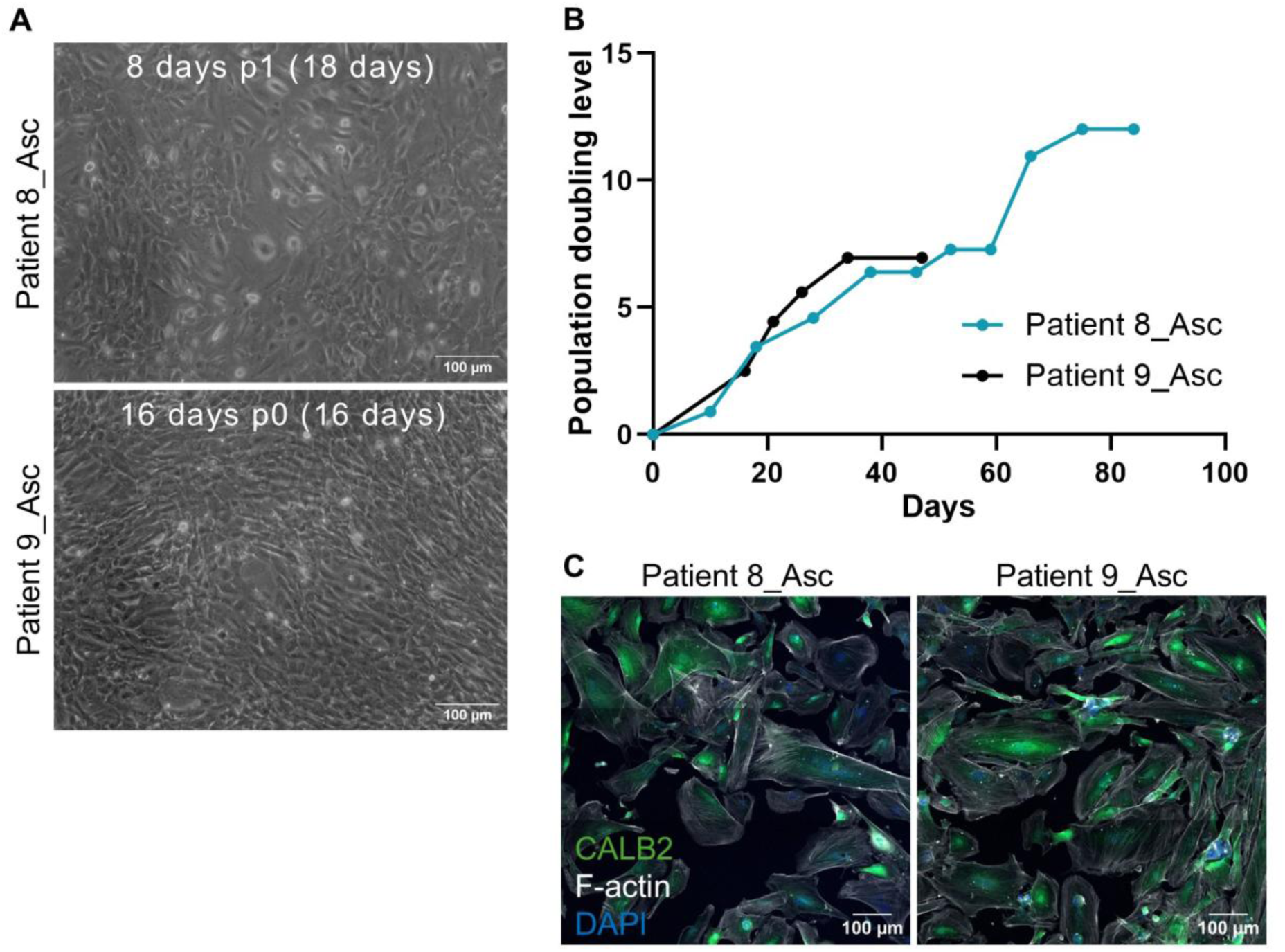
Mesothelial cell cultures. **A)** Representative 10X phase-contrast images of 2 patient-derived mesothelial cell cultures with the characteristic cobblestone morphology. The number of days in each passage, the passage number as well as the total number of days in culture are indicated in each image. Images were taken with a digital camera (Leica DFC340 FX) attached to an optical microscope. Scale bar: 100 µm. **B)** Chart displays the growth of mesothelial cell cultures as changes in population doubling level (PDL) over time. These cultures, which were established from cryopreserved enriched cell suspensions from ascites, displayed similar growth dynamics. PDL at each passage was calculated as described in Figure 2B. **C)** Representative 20X confocal fluorescence images of 2 patient-derived mesothelial cell cultures stained for CALB2 (i.e. calretinin, mesothelial cell marker) and F-actin (cytoskeleton). Images were taken with Zeiss Laser Scanning Confocal Microscope (LSM800) as 3 x 3 tiles. Scale bar: 100 µm. Abbreviations: Asc – ascites fluid. The data that was used to generate B) is provided in Supplementary Table 2. The single channel images from C) are provided as Supplementary Figure 3.

**Figure 4.**
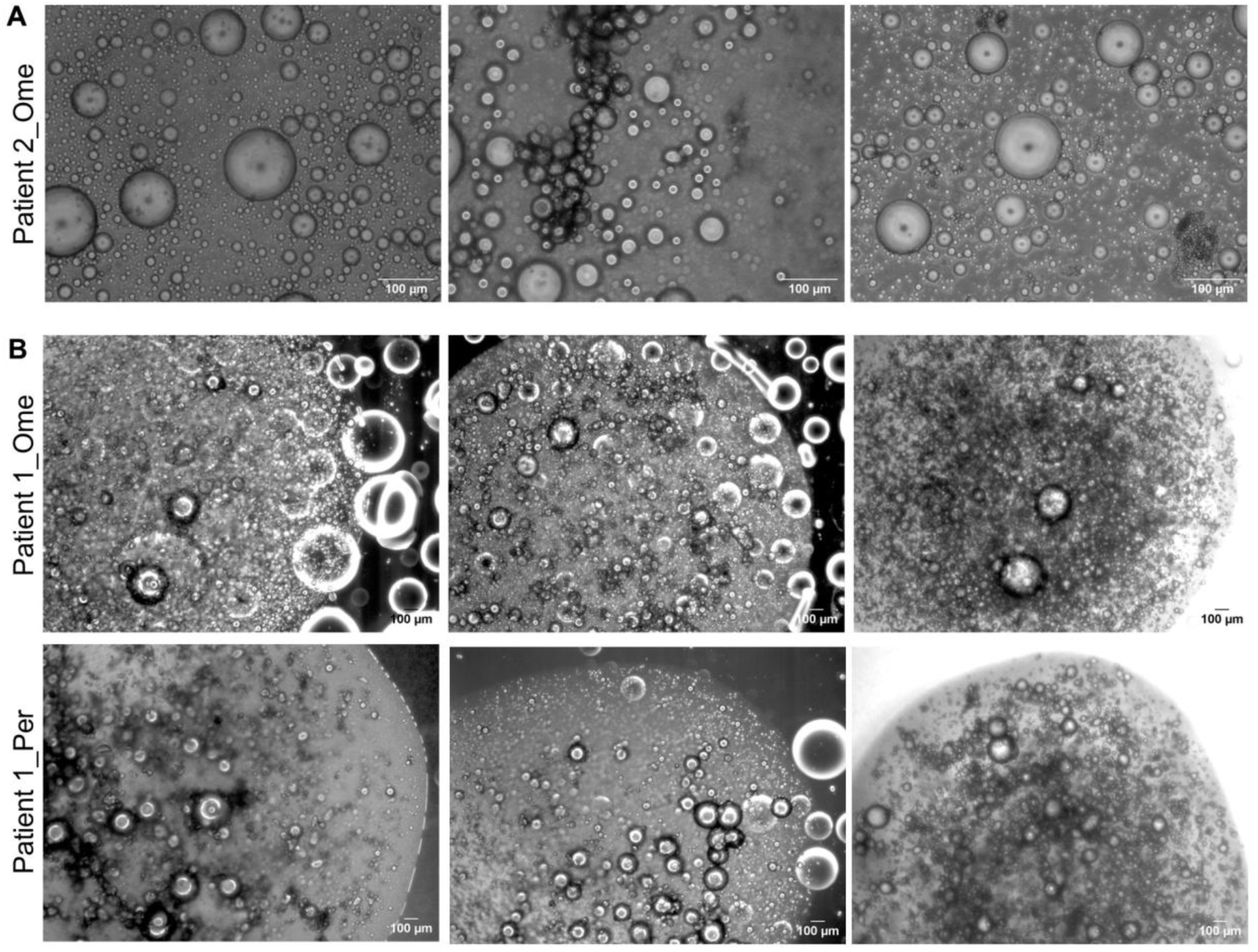
Adipocyte cultures. **A)** Representative 10X phase-contrast images of floating adipocytes and lipid vesicles after performing step 26. Images were taken with a digital camera (Leica DFC340 FX) attached to an optical microscope. Scale bar: 100 µm. **B)** Representative 2.5X phase-contrast images of adipocyte gels derived from 2 different tissues of the same patient (i.e. omentum – top row, peritoneum – bottom row) after performing step 33 (2 images on the left side). Images on the right show the adipocyte gels after 9 days of culture. Images were taken with a digital camera (Leica DFC340 FX) attached to an optical microscope. Scale bar: 100 µm. Abbreviations: Ome – omentum, Per – peritoneum.

The MC model described in this protocol can be generated in different formats (i.e. plate types) and used in various applications. In addition, by adding mCherry-labelled organoids (optional step), cancer cells can be easily separated from the stromal cells when imaged with a fluorescent microscope (**Figure 5**). The miniaturized MC model can be used to test patient drug responses to chemotherapy (and other drugs) in a high throughput manner (**Figure 6**), becoming a valuable tool for precision cancer medicine.

**Figure 5.**
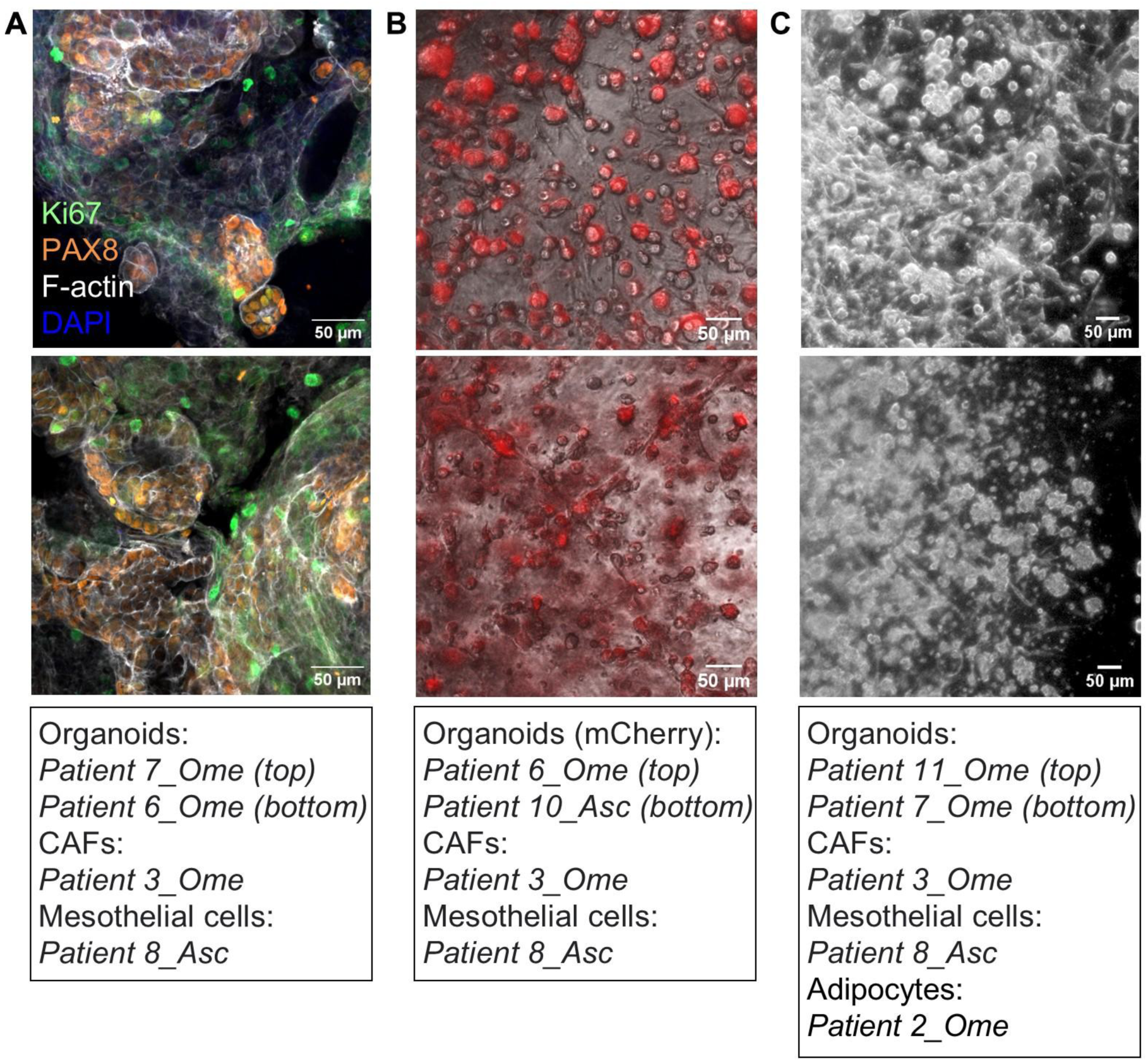
The multicellular culture model. **A)** Representative 20X confocal fluorescence images of the MC model. The model was grown in 20 µL matrix domes in a 6-well plate for 10 days, fixed and stained for Ki67 (growth marker), PAX8 (HGSC cancer cell marker) and F-actin (cytoskeleton). Images were taken with a Zeiss Laser Scanning Confocal Microscope (LSM800). Scale bar: 50 µm. **B)** Zoomed in insets from representative 4X merged phase-contrast and fluorescent images of the MC model. The model was grown in 10 µL of the mixture of matrices in a 384-well plate and imaged after 6 days with a digital microscope (EVOS M3000 Imaging System, Thermo Fisher Scientific). Scale bar: 50 µm. **C)** Zoomed in insets from representative 2.5X phase-contrast images of the MC model. The model was grown in 20 µL domes in a 6-well plate, exposed to an 8-hour chemotherapy treatment (10 µM carboplatin and 15 nM paclitaxel) on day 6, and used on day 12 to perform scRNA-seq. Images were taken on day 6 with a digital camera (Leica DFC340 FX) attached to an optical microscope. Scale bar: 50 µm. The origin of the cells used to generate each MC model is indicated below the images. The single channel images from A) are provided as Supplementary Figure 4.

**Figure 6.**
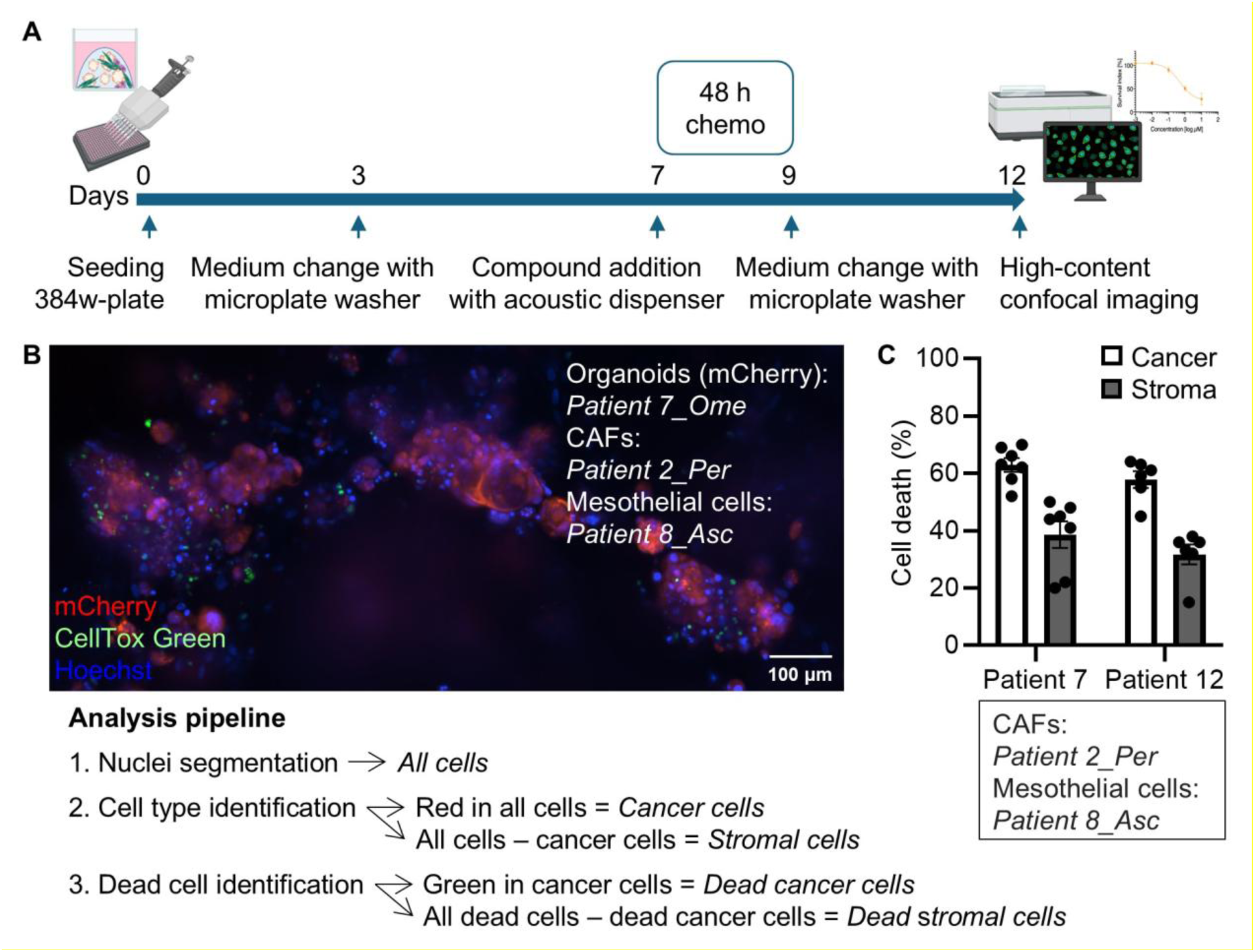
Example application of the MC model. **A)** Schematic of a high throughput drug sensitivity testing experiment with the MC model according to steps 69-82 of the protocol. **B)** Zoomed in inset from a representative 10X confocal fluorescence image of the MC model (the origin of the cells is indicated in the image). The model was grown in 10 µL of the mixture of matrices in a 384-well plate, handled for high throughput drug sensitivity testing as depicted in A, and imaged after 12 days with ImageXpress Confocal HT.ai high-content imaging system (Molecular Devices). The obtained images were analyzed with MetaXpress software (Molecular Devices) following the pipeline indicated below the image. Scale bar: 100 µm. **C)** Chart displays the percentage of dead cancer and stromal cells of the MC model (the origin of the cells is indicated in and below the chart). High throughput drug sensitivity testing was performed as depicted in A (including a 48-hour chemotherapy treatment with 50 µM carboplatin and 15 nM paclitaxel) and image analysis as indicated in B. Data are shown as mean ± SEM of triplicate wells and are representative of three experiments. The data that was used to generate C) is provided in Supplementary Table 3.

In our hands, processing the MC model for scRNA-seq as described in this protocol resulted in cell viability between 72-95% for 4 different patient-based models that had been left untreated or treated with carboplatin (10 µM) and paclitaxel (15 nM) for 8 h.

## Limitations

Establishing organoid and stromal cell cultures from patient material is an empirical process for which we cannot predict success. Therefore, it may not be possible to obtain all the different cell types required to generate the MC model from one single patient or sample type. Nevertheless, considering that the interpatient tumor heterogeneity in HGSC is mostly driven by cancer cells rather than the stroma, as evidenced by multiple single cells studies^5,6^, we deem that using patient unmatched stroma to generate the MC model is acceptable.

## Troubleshooting

### Problem 1

When adjusting the pH of collagen, too much NaOH is added, resulting in a pH above 8 and its polymerization before use (related to step 25 of Preparation of 2.25 mg/mL Type I collagen and step 63 of Generating the MC model).

### Potential solution

If the pH is just above the desired one, it may be possible to use the collagen after vortexing the tube with collagen shortly multiple times and placing it on ice for 1 min. Check the viscosity by taking up a small fraction with a pipette and assess the pH again. If the problem persists, take a new aliquot of collagen and perform the pH adjustment again. Remember that it is critical to add NaOH in small volumes and check the pH after each addition, particularly when the pH is close to 6-7.

### Problem 2

After centrifuging the fatty layer, it is not possible to clearly distinguish and separate the oily fraction and residual medium from the adipocyte fraction (related to step 7 of Handling fresh solid tumor samples).

### Potential solution

Proceed directly to step 25 (<u>Establishing adipocyte cultures</u>) and place the whole solution in culture. As adipocytes grow in suspension while other stromal cells attach to the cell culture surface, the cells will be naturally separated after being in culture for some days. In addition, red blood cells and immune cells will die because of the specialized Adipocyte medium.

### Problem 3

The adipocyte gels collapse and dissolve upon addition of medium (related to step 33 of Establishing adipocyte cultures).

### Potential solution

Collect the cells and medium into a tube, centrifuge at 200 x g for 5 min and aspirate the supernatant. Resuspend the cells in 5 mL of Adipocyte medium and repeat the centrifugation step. Make sure to manually remove all the medium using a pipette instead of automatic suction. Proceed to step 29 (<u>Establishing adipocyte cultures</u>).

### Problem 4

The stromal cultures are not phenotypically homogenous, and the expression of cell type-specific markers is low (related to step 20 of Establishing CAF cultures from tissue pieces and step 54 of Establishing mesothelial cell cultures from cryopreserved patient samples).

### Potential solution

Consider the possibility of enriching the culture for the cell population of interest with magnetic cell separation or flow cytometry cell sorting approaches.

### Problem 5

The mixture of matrices is very viscous and thoroughly resuspending the cell pellet and/or seeding the mixture in the 384-well plate before its polymerization is difficult (related to steps 63-65 of Generating the MC model and steps 71-73 of Miniaturizing the MC model for high throughput drug sensitivity testing).

### Potential solution

Make sure to keep all the reagents and materials as well as the cell pellet resuspended in the mixture of matrices on ice to slow down the process of matrix polymerization, and work at a steady pace. If needed, use wide bore pipette tips (alternatively cut the edge of the tips with sterile scissors) to facilitate handling of the viscous matrices.

## Resource availability

### Lead contact

Further information and requests for resources and reagents should be directed to and will be fulfilled by the lead contact, Lidia Moyano-Galceran.

### Technical contact

Technical questions on executing this protocol should be directed to and will be answered by the technical contact, Lidia Moyano-Galceran.

### Materials availability

This study did not generate new unique reagents. Previously established patient-derived organoids (as described in Senkowski *et al*.^1^) can be obtained through the OvaCure Collection and Auria Biobank https://www.ovacurecollection.com/

### Data and code availability

Original/source data are available as supplementary tables and figures.

## Acknowledgments

This work was supported by European Union’s Horizon 2020 research and innovation program under grant agreement no. 101063359 for CROC (LM-G). Views and opinions expressed are those of the author(s) only and do not necessarily reflect those of the European Union or the European Research Executive Agency (REA). Neither the European Union nor the REA can be held responsible for them. Funding was also received from Danish Cancer Society grant no. R374-A22457 (KW), Innovation Fund Denmark/ERA PerMed JTC2020 grant 0204-00005B (KW) and Novo Nordisk Foundation grant no. NNF21OC0070381 (KW).

Access to HGSC patient material was kindly provided through collaborations with: i) Prof. Sampsa Hautaniemi, University of Helsinki, and Dr. Johanna Hynninen, Turku University Hospital, and the European Union’s Horizon 2020 research and innovation program project DECIDER (grant agreement no. 965193), and ii) Dr. Sahar Salehi, Karolinska University Hospital, and the IPLA-OVCA clinical trial (trial no. NCT04065009).

We thank the staff of BRIC’s High-Content CRISPR Screens Facility and BRIC’s Light Microscope Facility for providing research infrastructure (Novo Nordisk Foundation Infrastructure grant number NNF20OC0061734) and excellent technical help. Plasmid pLEX307_SmaI-T2A-mCherry was a kind gift from BiOrigin ApS. Graphical abstract and Figure 6A were created with BioRender.

## Author contributions

**KW**: Resources, Funding acquisition; **DB**: Methodology; **LG-M**: Methodology; **WS**: Methodology, Resources; **LM-G**: Conceptualization, Methodology, Investigation, Formal analysis, Visualization, Writing - Original Draft, Project administration, Funding acquisition. All authors reviewed and approved the final manuscript.

## Supplementary Files

**Supplementary Figure 1. Uncropped immunoblots related to Figure 2C**.

**Supplementary Figure 2. Single channel images related to Figure 2D**.

**Supplementary Figure 3. Single channel images related to Figure 3C**.

**Supplementary Figure 4. Single channel images related to Figure 5A**.

**Supplementary Table 1. CAF growth data used in Figure 2B**.

**Supplementary Table 2. Mesothelial cell growth data used in Figure 3B**.

**Supplementary Table 3. Cell death data from the MC model used in Figure 6C**.

