## Supplementary figures and images for "Generation of an in vitro 3D multicellular culture model of ovarian high-grade serous carcinoma"

### Supplementary Figures

**Supplementary Figure 1**

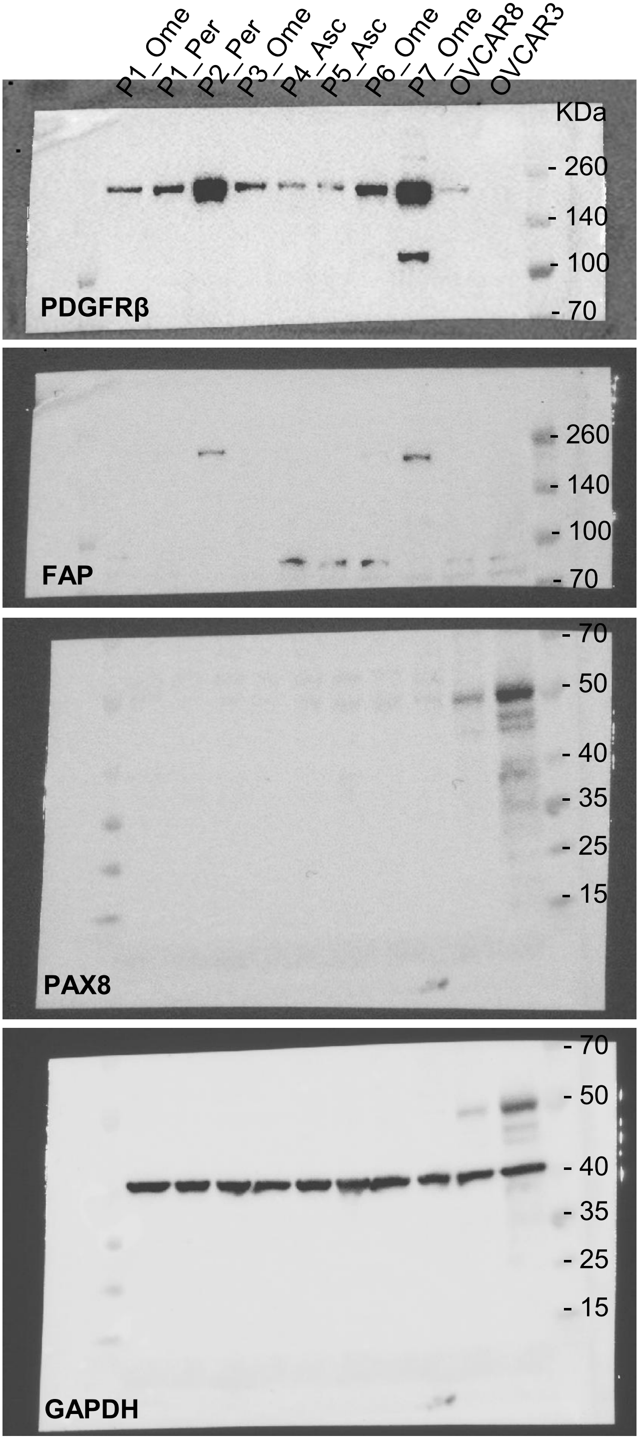

**Supplementary Figure 2**

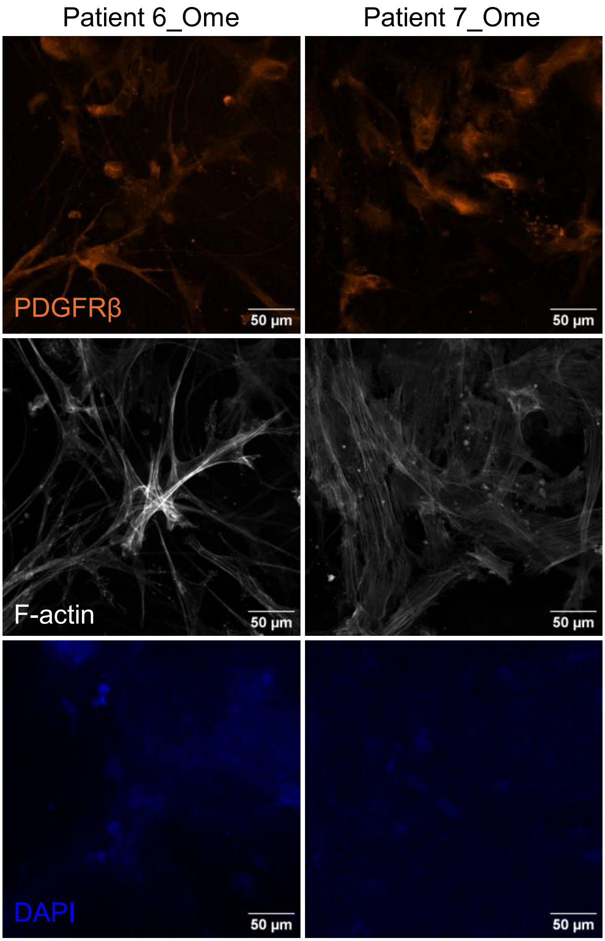

**Supplementary Figure 3**

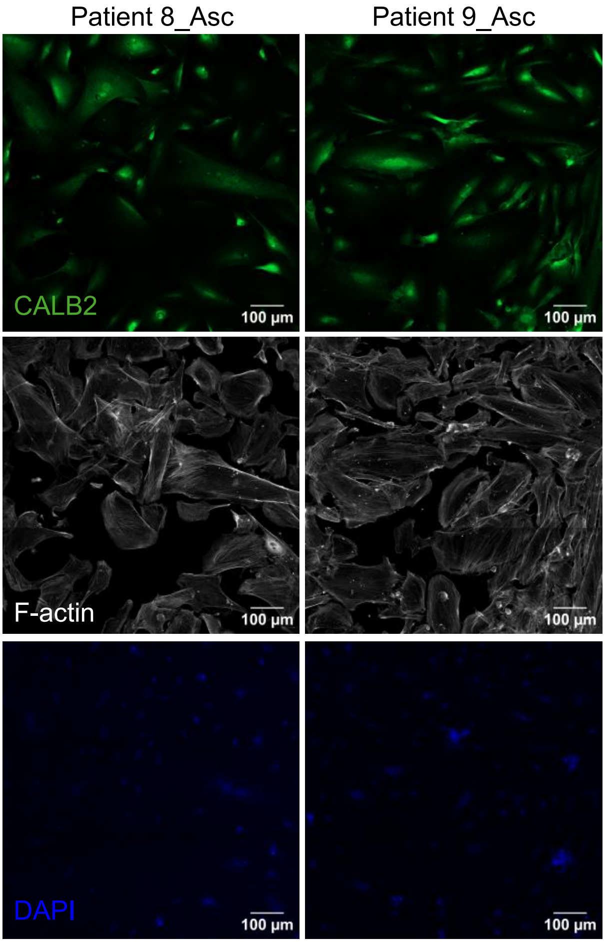

**Supplementary Figure 4**

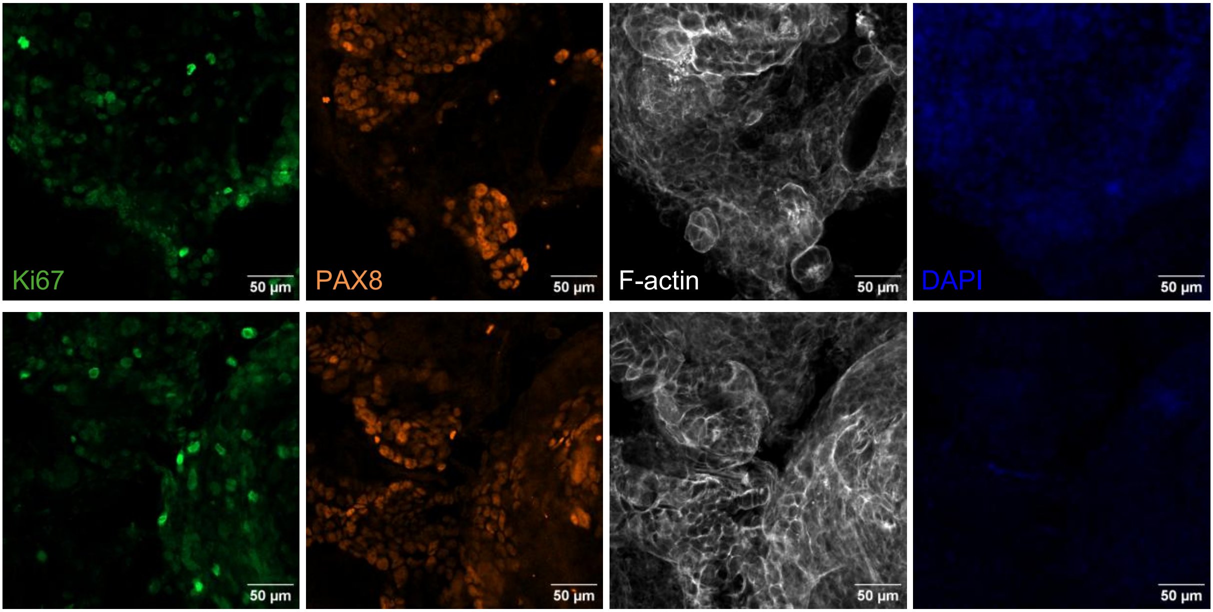
